# Animal Model Prescreening: Pre-exposure to SARS-CoV-2 impacts responses in the NHP model

**DOI:** 10.1101/2020.07.06.189803

**Authors:** Keersten M. Ricks, Andrew S. Herbert, Jeffrey W. Koehler, Paul A. Kuehnert, Tamara L. Clements, Charles J. Shoemaker, Ana I. Kuehne, Cecilia M. O’Brien, Susan R. Coyne, Korey L. Delp, Kristen S. Akers, John M. Dye, Jay W. Hooper, Jeffrey M. Smith, Jeffrey R. Kugelman, Brett F. Beitzel, Kathleen M. Gibson, Sara C. Johnston, Timothy D. Minogue

## Abstract

COVID-19 presents herculean challenges to research and scientific communities for producing diagnostic and treatment solutions. Any return to normalcy requires rapid development of countermeasures, with animal models serving as a critical tool in testing vaccines and therapeutics. Animal disease status and potential COVID-19 exposure prior to study execution may severely bias efficacy testing. We developed a toolbox of immunological and molecular tests to monitor countermeasure impact on disease outcome and evaluate pre-challenge COVID-19 status. Assay application showed critical necessity for animal pre-screening. Specifically, real-time PCR results documented pre-exposure of an African Green Monkey prior to SARS-CoV-2 challenge with sequence confirmation as a community-acquired exposure. Longitudinal monitoring of nasopharyngeal swabs and serum showed pre-exposure impacted both viral disease course and resulting immunological response. This study demonstrates utility in a comprehensive pre-screening strategy for animal models, which captured the first documented case of community-acquired, non-human primate infection.

**One Sentence Summary:** Pre-exposure to SARS-CoV-2 affects biomarker responses in animal models, highlighting a need for robust pre-screening protocols prior to medical countermeasure studies.

---

Longstanding awareness of coronavirus as an infectious disease, from the “seasonal-cold” to the smaller outbreak clusters of severe acute respiratory syndrome (SARS1) and Middle Eastern respiratory syndrome (MERS), informed the infectious disease and public health community of the potential for the current outbreak(*1-4*). One-Health efforts, led by multi-organizational teams such as PREDICT, highlighted a wide range of novel coronaviruses circulating in animal reservoirs and the human-environmental interface as possible outbreak candidates(*5, 6*). Previous Ebola virus outbreaks in West Africa and DRC invigorated discussions on pandemic preparedness and rapid response to “Disease X”; however, the globe faced a stark reality in late 2019, with the ongoing SARS-CoV-2 pandemic exposing critical gaps in the ability to respond(*7, 8*). Researchers are moving at a rapid pace to fill these gaps with development of diagnostics, vaccines and therapeutics(*9, 10*). For these efforts to succeed and impact the spread of the virus, there is the critical need for well-characterized assays to identify the virus, assess the immune response post-exposure, and determine efficacy of vaccines and therapeutics(*11*). Despite extensive research and animal modeling post-SARS-CoV-1 and MERS outbreaks as well as the rapid rate of research on SARS-CoV-2, little is known regarding longevity of immune protection, vaccine efficacy, or reinfection potential for this new coronavirus(*12-17*).

Globally, vaccine and therapeutic groups are addressing the need for a better understanding of pathogenesis and appropriate animal models by tackling specific questions such as non-human primate species selection, potential for viral reinfection, and virus detection and immunological profiling(*18-22*). Specifically, Haagmans et.al found no clinical signs of disease in a cynomolgus macaque challenge model but noticed viral secretions in the nose and throat of challenged monkeys with disease severity to be between that of SARS1 and MERS(*23*). In a rhesus macaque reinfection model by Qin et.al, NHPs challenged post-recovery did not exhibit any evidence of re-infection with SARS-CoV-2(*24*). In addition, many groups are detecting virus and subsequent immune response in asymptomatic, mild, and severe COVID patients(*25-27*). Data from these studies suggest a differential response in both the species selected to initial SARS-CoV-2 versus a reinfection models. This also impacts the diagnostic context where unanswered questions regarding concordance between immunological data and correlates of protection inhibit efforts for instituting “Immunity passports”(*28*). Diagnostic assay development and validation efforts focus on diagnosis and post-exposure screening of the human population, however, application of a diagnostic toolbox across animal models is essential for therapeutic and vaccine efforts for a complete understanding of disease progression and treatment efficacy(*29, 30*).

Diagnostic and animal model assays generally parse into two bins based on target biomolecule: nucleic acid or protein diagnostics. Both types of assays, in a qualitative and quantitative context, inform animal model and countermeasure development (Table S1). For instance, real-time PCR is used for both pathogen detection (qualitative) and determining viral load (quantitative). Pathogen-specific IgM and IgG antibody immunological assays characterize the infection-specific host response and, diagnostically, are commonly used to screen for recent or previous pathogen exposure. Critical to the countermeasure development is the understanding of the protective response generated through exposure. Characterizing this immune response using microneutralization and PRNT assays adds specificity and qualitative data to immune response. Coupled with existing animal model data, animals used for COVID-19 model and countermeasure development should be screened for current and previous exposure to the study organism given the significant impact previous exposure can have on the study results.

High transmissibility and prevalence of this novel coronavirus presents a unique challenge to animal model developers. Pre-screening animals for past exposure against most emerging infectious diseases, such as viral hemorrhagic fevers and other lesser-known arthropod borne viruses, often only requires a simple ELISA. In these NHP cohorts, the risk of prior exposure or having cross-reactive antibodies against the challenge agent is low. Here, we demonstrate the need for a fully developed molecular and immunological toolbox prior to animal challenge when considering animal models against a disease agent of pandemic potential. We present an integrated assay development approach (PCR, sequencing, immunoassays, and neutralization assays) for a SARS-CoV-2 species selection study. In the prescreen application of this toolbox, an African Green (AGM) non-human primate (NHP) showed evidence for previous exposure to SARS-CoV-2 prior to challenge with the Institute stock (Fig 1). Additionally, use of this integrated screening approach post-challenge showed the immunological and viral load were significantly impacted by this pre-exposure across longitudinal assessments.

**Fig 1.**
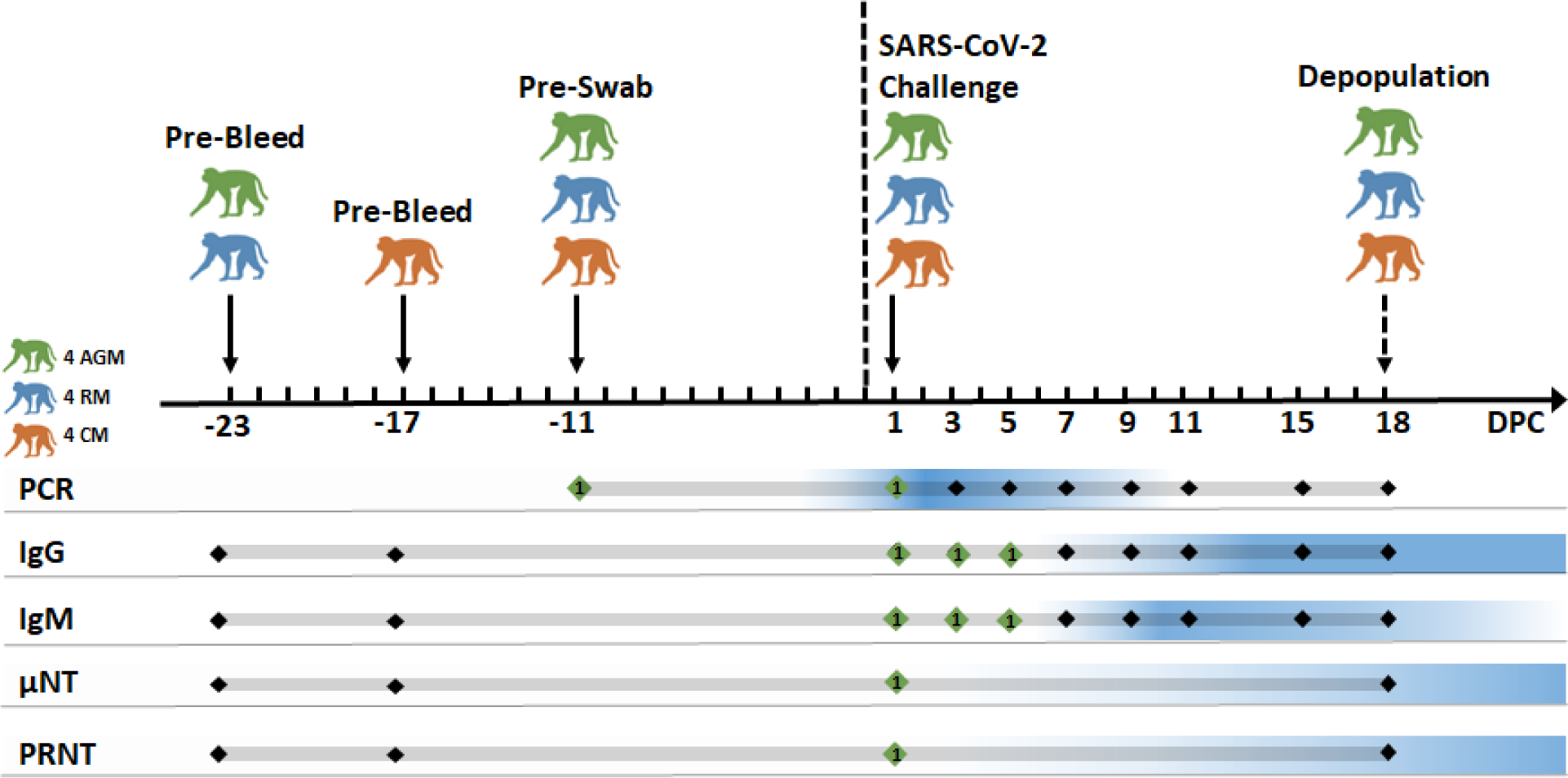
Summary of pre-challenge and post-challenge sample analysis and results. Green diamonds indicate a positive assay result for the pre-exposed AGM NHP, where the remaining NHPs were negative. Blue gradient indicates where the average response of the NHPs was positive for each assay.

## Results

### Immunoassay development

We took a multi-pronged assay development/implementation approach in order to mitigate risk prior to animal model development by producing in-house assays as well as integrating a commercial, EUA approved, ELISA kit. Commercial sources and academic collaborators provided SARS-CoV-2 recombinants to develop a multiplex, bead-based immunoassay on the Magpix platform. Our previous efforts used this method to qualitatively detect both IgM and IgG response, against glycoprotein and nucleoprotein targets, in animal models, such as CCHFV, RVFV, and VEEV, thus simplifying platform selection, assay optimization, and matrix testing, as we could leverage our successes in utilizing this platform for longitudinal biomarker assessment in serum (*31, 32*). The newly developed multiplex Magpix assay targeted full trimeric spike protein, S1 subunit, receptor-binding domain, nucleoprotein, and SARS1 spike protein (Fig S1). We ordered the SARS-CoV-2 S1 IgG ELISA kit from Euroimmun, to run in parallel to our in-house Magpix multiplex, as there was substantial validation data for the assay.

### Real-time RT-PCR development

We optimized two real-time, qualitative, RT-PCR assays, (Q)N2 and GP. These assays used different reagents than the Center for Disease Control (CDC) Emergency Use Authorization (EUA) reagents due to supply shortages, thus establishing an independent supply chain from human EUA testing. The two assays showed similar performance when tested against serially diluted SARS-CoV-2 nucleic acid, with a slight increase in sensitivity with the (Q)N2 assay compared to the GP assay (Fig S2A and 2B). We observed no cross-reactivity across a 103-organism exclusivity panel, including eight different coronaviruses. Clinical matrices contain inhibitors that can negatively impact nucleic acid extraction and real-time PCR performance. Matrix testing with the more sensitive (Q)N2 assays showed similar performance in both PBS and NP swab matrix at higher virus concentrations; however the LOD was lower in less complex PBS matrix (Fig S2C). Sixty replicate extractions with singlet (Q)N2 testing resulted in all replicates testing positive at a virus concentration of 0.614 PFU/ml (Fig S2D).

### SARS-CoV-2 Microneutralization assay development and optimization

Leveraging past successes developing microneutralization assays, we optimized a rapid, high-throughput method for evaluating therapeutic countermeasures against live SARS-CoV-2 virus. Screening of four commercially available antibodies identified a SARS-CoV-1 NP-specific cross-reactive antibody capable of detecting SARS-CoV-2 infected cells in an immunofluorescence-based assay (IFA). Evaluation of mean fluorescent intensity of this serially diluted antibody on uninfected and SARS-CoV-2 infected cells determined the optimal immunostaining concentration. Peak intensity began to decrease near 1:4500 dilution of primary antibody resulting in a 1:5000 dilution of primary antibody for all subsequent immunostaining. (Fig S3A). Vero-E6 cells were equally, if not more, permissive for SARS-CoV-2 infection compared to Vero-76 cells inoculated with an equivalent multiplicity of infection (MOI) (Fig S3B). At 24 hours, percent infection ranged from 30% at MOI 0.6 to less than 2% at MOI 0.007 with standard deviation between 0.6 and 2.4 (Fig S3C). By 48 hours, MOIs between 0.6 and 0.022 resulted about 60% infection with standard deviations between 1.7 and 2.7. Percent infection at MOI 0.007 resulted in about 40% infection and standard deviation of 8.5%.

The primary focus for this effort was developing an assay to evaluate virus entry inhibition, therefore, we opted for a 24 hour assay in order to limit rounds of virus replication. Across several independent experiments with SARS-CoV-2/MT020880.1 at an MOI of 0.4 percentage of infected cells within assays were consistent with intra-assay coefficient of variance (COV) ranging from 4.9% to 6.8% and 2.7% to 9.5% based on the viral stock used (Fig S3D). Similarly, mean percent infection was also consistent for independent stocks (mean = 43.2%) and (mean = 57%) with inter-assay COV of 15.6% and 14.2%, respectively (Fig S 3E). Data compiled from independent experiments with cells infected with SARS-CoV-2 at an MOI of 0.2 or 0.08 showed intra-assay variability across replicate wells was more consistent at MOI 0.2 as compared to with MOI 0.08 (Fig S3F).. Based on these results, we performed all subsequent experiments at 24-hour assay and an MOI 0.2.

### Pre-study NHP Baseline Screening

Prior to the study start, we screened serum from 12 total NHPs (4 AGM, 4 RM, 4 CM) against the Euroimmun ELISA, COVID Magpix Multiplex, µNT, and PRNT. Study coordinators took pre-bleed samples between 23 and 17 days pre-challenge. All pre-bleed serum samples were negative by immunoassay and Plaque Reduction Neutralization Test (PRNT) assays (Table S2). As part of re-iterative prescreening for COVID, study coordinators also took pre-study NP swabs from these NHPs 11 days pre-challenge (Fig 1) Surprisingly, the left nare NP swab of one AGM tested positive by the (Q)N2 assay and subsequently by the GP assay (Table S3). Sequencing of the extracted nucleic acid material showed >95% coverage to the SARS-CoV-2 reference genome and found to be in a separate clade than the USAMRIID challenge stock material. Institution stocks are based on the WA1 (MT020880.1); however, the AGM swab was phylogenetically closer to isolates that have been circulating in the US and Europe (Fig S4). The study team swabbed all NHPs again, including the positive AGM, immediately prior to challenge. The left nare NP swab was positive with both assays, and almost half of the right nare NP replicates tested positive with the GP assay. No other NHP in the study was PCR positive by NP swab prior to study start (Fig 1).

### Longitudinal Assessment of Immune Response and Viral Load

We screened all time points post-challenge by ELISA, MAGPIX, and PCR assays. All NHPs showed no IgG response on Day 1 post-challenge (PC), with the exception of the previously exposed AGM, AGM1 (Fig S5A-C). Measurable IgG response in the Euroimmun ELISA started at Day 11 PC and increased through to terminal bleed. Interestingly, the previously exposed AGM IgG response plateaued at Day 7 PC; however, we observed an anamnestic response at Day 11 PC (Fig 2A). Further analysis of this immune response using the COVID IgM and IgG MAGPIX multiplex immunoassay at all days PC showed specific IgM signal only to SARS2 glycoprotein antigens, as these assays yielded no measurable signal for the SARS1 spike or the SARS2 nucleoprotein (Fig 2B, Fig S6). IgM signal was detectable above baseline for the average of each NHP species by Day 9 for the AGM and RM and Day 7 for the CM, for all three glycoprotein derivatives. IgG signal was also detectable above baseline at Day 9 for AGM and RM and Day 7 for CM (Fig S7). The Magpix immunoassay showed higher relative sensitivity than the Euroimmun ELISA kit, with detection of IgG signal in the NHPs 2-4 days sooner. Based on this assay, there was no observed restimulation in the IgM response of the pre-exposed AGM post-challenge, as noted for IgG (Fig 2B). IgG response from the pre-exposed AGM was statistically significant from the 15 other NHPs at the day of challenge, however, there was no statistical difference serologically between that AGM and the rest of the cohort at terminal bleed (Fig S8). In screening the NP swabs by PCR, the cohorts had detectable nucleic acid starting at D3 through D9-11, with one NHP in each species cohort having detectable nucleic acid by NP swab on the day of termination (Fig S5D-F). Notably, AGM1 had measurable nucleic acid on D1 through terminal bleed at lower relative levels of viral load exhibited by other members of the cohort post challenge (Fig 2C).

**Fig 2.**
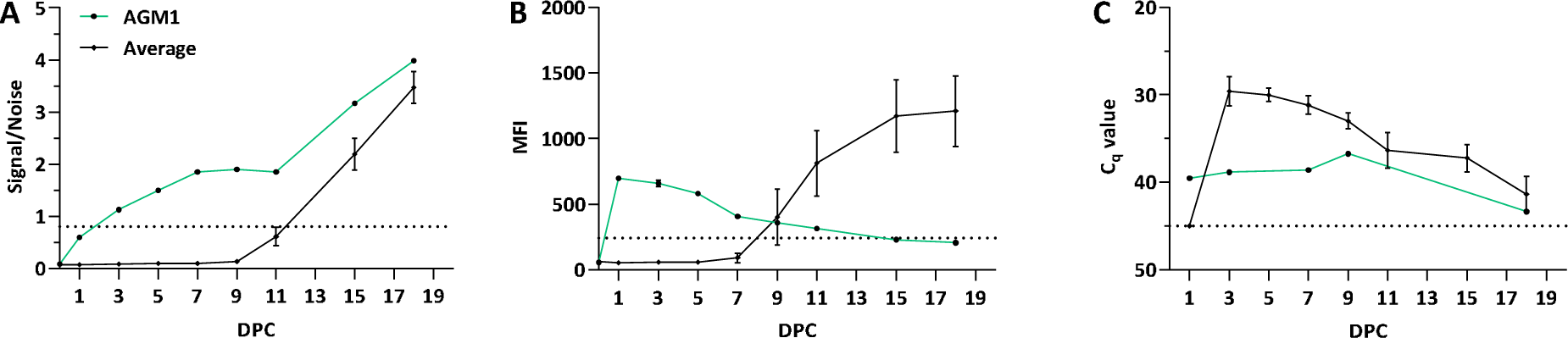
Longitudinal Immune and Viral Response. Longitudinal response for A) IgG, B) IgM, and C) PCR for the pre-exposed AGM, AGM1, as compared to the average response of the remaining study NHPs. Longitudinal samples were collected on day 1 and every two days thereafter. Subsequent sampling was done on days 15 and 18. Serum samples were screened by commercial Euroimmun SARS-CoV-2 S1 IgG ELISA kit or Magpix multiplex for IgM. The direct ELISA kit measures IgG response against the S1 subunit of the spike glycoprotein. The analogous S1 IgM assay from the Magpix multiplex is shown in panel B.

### Post-Challenge neutralization response

PRNTs and microneutralization assays demonstrated the neutralizing activity of serum collected at time of challenge and at the termination of the study (day 18 post-challenge). By PRNT, nearly all animals showed no neutralizing activity at time of challenge with PRNT80 titers less than 1:20 (Fig S9). The sole exception being the previously exposed AGM1, which had a PRNT titer of 1:160. In contrast, assessment by microneutralization assay demonstrated neutralizing activity of serum collected at time of challenge in all AGM serum samples with calculated half-maximal inhibitory concentration (IC_50_) from 1:212 to 1:1274 (Fig 3, Table S4). In agreement with ELISA and PRNT data, previously exposed AGM1 had the highest pre-challenge IC_50_. Neutralizing activity of RM and CM pre-challenge serum was similar to that of human negative control serum (Fig S10). On day 18 post-challenge, PRNT80 titers increased to between 1:160 and 1:2560 (Fig S9). Likewise, microneutralization assay IC_50_ increased 5-95 fold, the greatest fold difference occurring in RM and CMs (Table S4). Neutralizing titer in previously exposed AGM1 remained largely unchanged between time of challenge and day 18 in both assays. The fold-change in _log_IC50 microneutralization response in the average AGM cohort was noticeably higher than that of the pre-exposed AGM (Fig 3).

**Fig 3.**
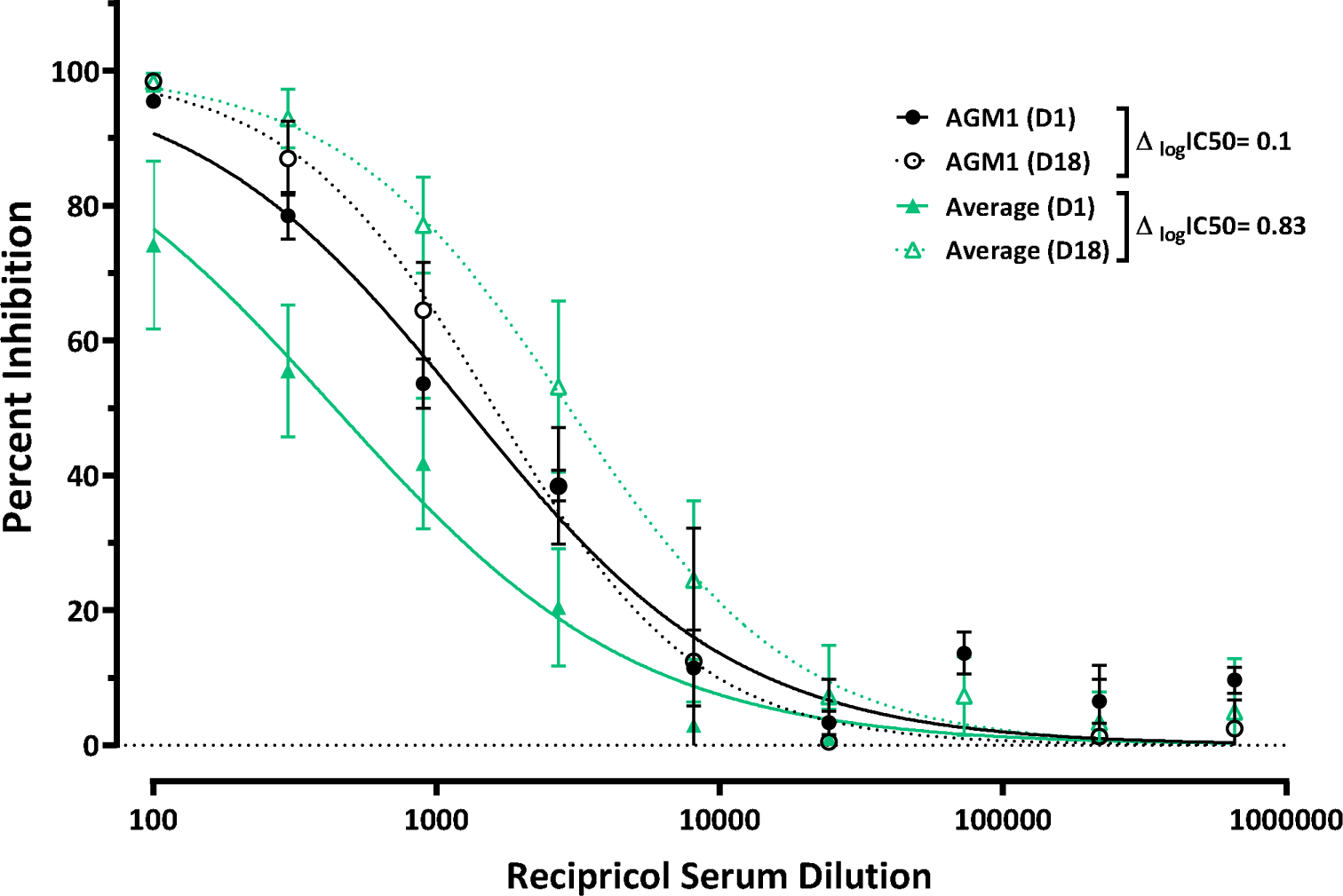
Neutralization activity of AGM NHP serum. Serially diluted serum (pre-challenge and day 18 post-challenge), mixed with pre-tittered SARS-CoV-2, was added to Vero-E6 cells at an MOI of 0.2. Cells were fixed 24 hours post-inoculation and immuno-stained with virus specific antibody to enumerate the number of infected cells by fluorescent microscopy (Operetta high-content imaging). Percent of infected cells was determined using Harmony software and percent inhibition calculated relative to untreated cells. The fold-change in the logIC50 of the pre-exposed AGM was relatively elevated in comparison to that of the average AGM response.

Mapping of all assay results showed the pre-exposure impact on neutralization and other relevant diagnostic targets (Fig 4). These data showed the dramatic increase from D1 to D18 in all diagnostic targets (RT-PCR, IgM and IgG) and neutralization (PRNT and microneutralization) across the entire cohort with the exception of the pre-infected cohort NHP AGM1. In NHP AGM1, we observed a generally static level of neutralization. IgG titers rose over the course of the study while, interestingly, IgM levels dropped (Fig 4, Fig S11).

**Fig 4.**
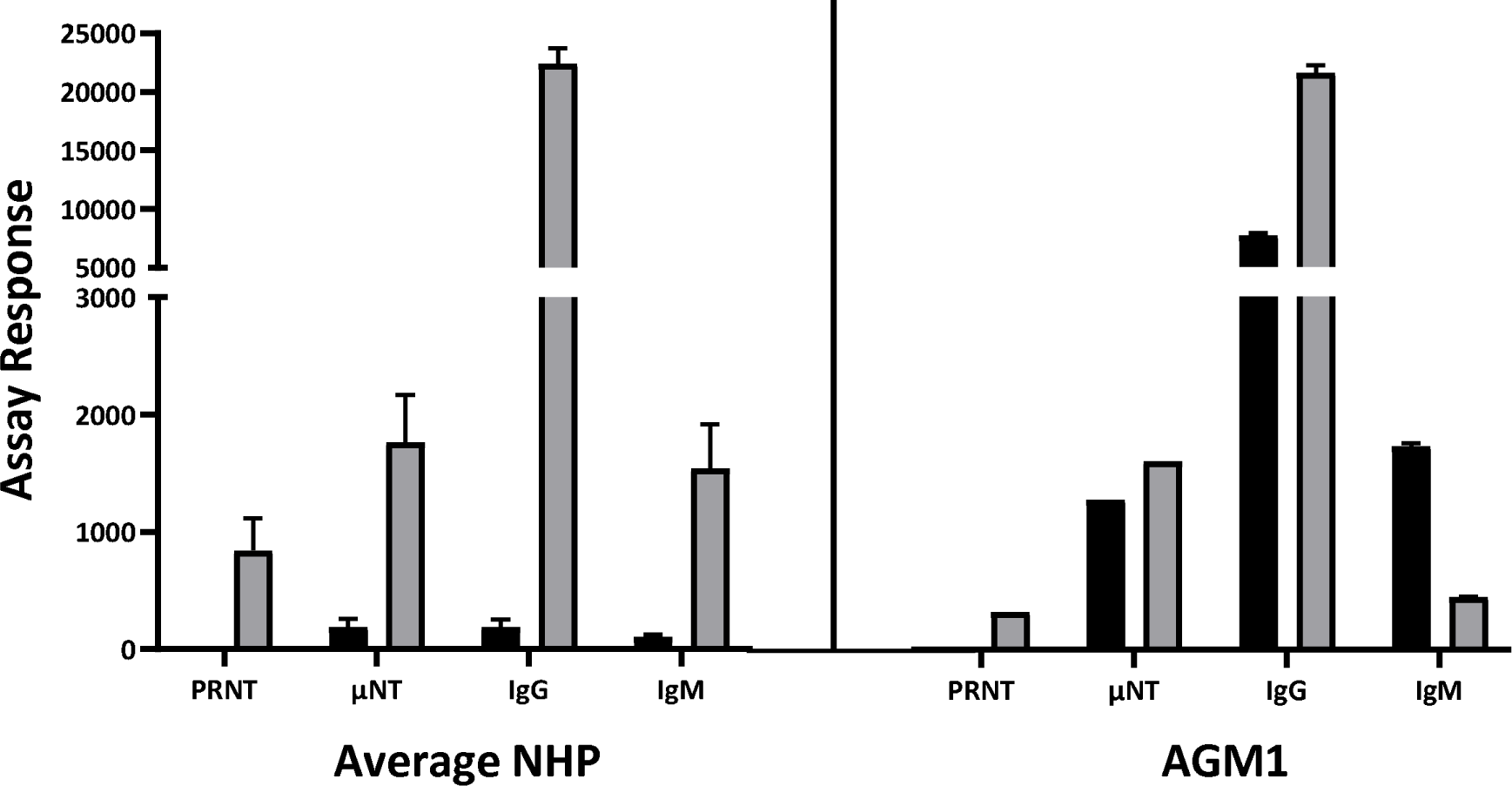
Pre-exposed NHP compared to Average NHP. Comparison of AGM1 to Average NHP response for each screening assay. The pre-exposed AGM generally had no consistent fold-change from D1 (gray bars) to D18 (black bars) whereas there was measurable positive fold-change for each assay for the average NHP cohort.

## Discussion

A key element to a modern society’s fortune in weathering disease outbreaks, and especially pandemics such as COVID-19, is its ability to coordinate an effective scientific and societal response. The earliest recorded pandemic, the Athenian Plague of 430 BC, killed more than 25% of the population of Athens during the Peloponnesian War and was theorized to have been caused by typhoid fever(*33*). In a more devastating fashion, 1918 Spanish Flu spread all over the world in the middle of World War I, killing more than 50 million and infecting more than 25% of the world’s population. Common to post-pandemic society after each event were the associated advancements made in public health response and infectious disease research(*8, 34, 35*). To mitigate the devastating effects of disease spread and rising mortality rates, there must be rapid response toward isolating the disease causing agent, diagnosing affected patients, treating those infected, managing further spread of the disease, and preventing future outbreaks(*36*). Rapid development of animal models is critical to response efforts in order to understand disease pathogenesis and serve as a platform for testing medical countermeasures.

The high transmissibility rate of COVID-19 and the dire need for medical countermeasures necessitates rapid initiation of animal studies to develop disease models for vaccine and therapeutic testing(*37*). Additionally, workflow and PPE considerations must be taken to mitigate pre-exposure risk of NHP colonies to fomites or other potentially infectious exposure routes. Pre-infection of the NHP in question occurred in spite of enhanced ABSL3 USAMRIID NHP handling procedures and represent a personal protective equipment (PPE) stance above and beyond what is currently being applied at other premier institutes. Previous SARS-CoV-1 and SARS-CoV-2 research shows distinct differences in host NHP response to challenge based on whether the animal is naïve or was pre-exposed to the virus(*24*). Therefore, we initiated the first documented, thorough pre-screening strategy prior to a species selection study to ensure the NHPs had no previous exposure markers. To do so required development of a COVID-19 diagnostic toolbox comprised of ELISA, PCR, multiplex IgM/IgG immunoassay, and neutralization assays to establish a full pre-challenge baseline and capture of post-challenge image of the SARS-CoV-2 animal model. Applying these tools in a prescreening context, we discovered one of the AGM NHPs had been previously exposed to SARS-CoV-2 with a community-acquired infection; the first documented case of community acquired NHP infection. Root cause analysis has not been undertaken, however; sequencing confirmed this was a different strain than the isolate used for challenge with sequence data showing this isolate to be of European and not Asian descent. Because this strain was not the a derivative of the challenge stock, this demonstrated potential for contamination of a naïve NHP cohort from the environment even when extraordinary preventive measures are taken to mitigate such risks.

Coupled with high transmissibility of the virus itself, the discovery of transmission from the community to an NHP has serious implications for future vaccine and therapeutic developmental efforts. All labs moving forward in SARS-CoV-2 countermeasure testing should consider pre-screening respective animal cohorts and implement measures to minimize the unintentional community to cohort virus transfer. Ultimately, responsibility of reporting community acquired NHP infections resides with United States Department of Agriculture (USDA). Reporting and better understanding of the breadth of community-acquired infection within NHP colonies will further demonstrate the impact and necessity for pre-screen cohorts prior to study initiation.

Analysis of the pre-exposed AGM to the cohort average post-challenge showed an inversion for all of the diagnostic markers (Fig 4). Detectable nucleic acid was present in the NP swab of AGM1 on D1 in addition to the presence of anti-SARS-CoV-2 IgG, IgM, and neutralizing antibodies. Interestingly, the relative level of viral load was present but consistently lower than the rest of the cohort and did not show marked increase across longitudinal evaluations. Additionally, IgG and IgM response in AGM1 were elevated on D1 relative to the baseline signal of the other NHPs. IgG appeared to be “re-stimulated” on D11 in concordance with the rest of the cohort; however IgM response slowly declined over the course of the study (Fig 2). Lastly, all of the NHPs had a net positive fold change in neutralizing antibody response from D1 to D18, while AGM1 had no significant net change (Fig 3). These data show the critical nature of a rapid but rational pre-screening response as the risk for not pre-screening prior to animal studies is evidenced in the alterations in these responses in pre-infected NHP. Together with the discovery of the pre-exposed NHP, this research raises questions regarding a larger discussion on the necessity of pre-screening in a mid-pandemic research community, implications for re-infection models, identification of true correlates of protection, and protective quality of future vaccines. These data clearly show that SARS-CoV-2 pre-exposure will have dramatic impact on any evaluation for vaccine or therapeutic efficacy and protective response due to countermeasure treatment.

Longitudinal data for COVID-19 diagnostic markers could be impactful for currently implemented diagnostic algorithms, the human screening for prior infection and potential for successful vaccine efforts(*38*). IgG and IgM co-induction, seen here, parallels current research in human cohorts with a non-phasic general increase in both immunoglobulins(*39, 40*). IgM is typically an early response indicator of the body’s adaptive immune response in the first week post-infection; however this response appears to be delayed for closely monitored SARS-CoV-2 infections. Generally, increase in IgG titers reflect neutralizing activity, a potential indicator of protection, thus giving some measure of confidence in establishing “immunity passports” for populations that seroconvert. However, recent studies showed not all those infected yield a productive or detectable immunological response(*28*). Additionally, evidence from the pre-infected NHP for both an anamnestic immunologic response and gross symptoms of productive SARS-CoV-2 infection suggest future vaccine efforts may have issues in producing durable, protective immunity. A SARS-CoV-2 reinfection model will need further investigation to establish whether data shown here gives evidence for a vaccine production design more similar to classic vaccination schedules or if it will require periodic updates similar to Flu. In general, further investigation will be required to establish what impact re-infection will have on COVID-19 diagnostic and countermeasures.

COVID-19 is not the first pandemic nor will be the last this generation will experience. Beyond the implications for prescreening animals prior to study, there are lessons in pandemic response that can be learned in time-to-product maturity and how we prioritize research efforts. Prioritizing assay development meets critical aspects for both diagnostic and countermeasure development needs. There will always be a need to develop diagnostics, vaccines and therapeutics in rapid response efforts; however, neglecting to clarify the relationships between diagnostic windows and matrices, therapeutic targets and toxicity, and vaccines and correlates of protection will only delay resolution of the COVID-19 crisis. To facilitate this resolution, we openly invite collaboration and can share more detailed protocols and development processes to stand-up these assays in other locations.

## Acknowledgements

The authors would like to thank the USAMRIID Comparative Medicine staff for all of their hard work in collecting NHP samples during the species-selection study to support this work.

## Author Contributions

**KR:** Conceptualization, methodology, formal analysis, writing, review & editing, visualization, funding acquisition. **AH:** Conceptualization, methodology, formal analysis, review & editing. **JK:** methodology, formal analysis, review & editing. **PK:** Investigation and formal analysis. **TC:** Investigation and formal analysis. **CJ:** Investigation and review & editing. AK: Investigation. **CO:** Investigation. SC: Investigation. **KD:** Investigation. **KA:** Investigation. **JD:** Conceptualization, supervision, funding acquisition. **JH:** Conceptualization, supervision. **JS:** Investigation. **JK:** Supervision. **BB:** Investigation. **KG:** Supervision. **SJ:** Resources. **TM:** Conceptualization, supervision, funding acquisition, writing, review & editing, project administration.

## Data and materials availability

All data is available in the main text or the supplementary materials.

