## Supplemental Materials for "Animal Model Prescreening: Pre-exposure to SARS-CoV-2 impacts responses in the NHP model"

**This PDF file includes:**

Methods

Figures. S1 to S11

Tables S1 to S4

Methods

Ethics Statement

This work was supported by an approved USAMRIID Institutional Animal Care and Use Committee (IACUC) animal research protocol in compliance with the Animal Welfare Act, PHS Policy, and other Federal statutes and regulations relating to animals and experiments involving animals, AP-20-009. The facility where this research was conducted is accredited by the Association for Assessment and Accreditation of Laboratory Animal Care, International and adheres to principles stated in the Guide for the Care and Use of Laboratory Animals, National Research Council, 2011. Approved USAMRIID animal research protocols undergo an annual review every year.

Generation of authentic SARS-CoV-2 stock

Vero-76 cells were inoculated with SARS-CoV-2/MT020880.1 at a multiplicity of infection (MOI) = 0.01 and incubated at 37°C/5% CO2/80% humidity. At 50 hours post-infection, cells were frozen at -80°C for 1 hour (hr), allowed to thaw at room temperature, and supernatants were collected and clarified by centrifugation at ~2500xg for 10 minutes. Clarified supernatant was aliquoted and stored at -80°C. Sequencing data from this virus stock indicated a single mutation in the spike glycoprotein (H655Y) relative to Washington state isolate MT020880.1.

NHP Study Sample Collection and Processing

As part of a pre-screening procedure for evidence of active or past SARS-CoV-2 infection, the three cohorts of NHPs to be used in a species down-selection study for COVID-19, African Green Monkeys (AGM), Cynomolgus macaques (CM), and Rhesus macaques (RM) were anesthetized between 23 and 17 days before challenge for blood collection, and again 11 days prior to challenge for nasopharyngeal (NP) swab collection. A final combined blood and NP swab collection was conducted immediately prior to challenge (Day 1). Post-challenge serum and NP swabs were collected at indicated time points and a final terminal bleed was conducted 18 days post-challenge. Whole blood was collected from a peripheral vein and stored in K3 EDTA tubes followed by serum isolation using a gel-based serum separator (Sarstedt, Numbrecht, Germany) and storage at -80°C for subsequent serological analysis. Individual nasopharyngeal swabs were collected from each NHP nare using a BBL CultureSwab with a rayon soft swab tip. After collection, swabs were deposited in Viral Transport Media (1x Hanks Balanced Salt Solution supplemented with 2% Fetal Bovine Serum, 100 μg /mL Gentamicin, and 0.5 μg /mL Amphotericin B). Samples were then vortexed, swabs removed, and 100 uL of sample was combined with 300 ul TRIzol LS to inactivate any virus present, and frozen until subsequent extraction and RT-PCR analysis.

Real-time RT-PCR

*Assay Design.* Two different real-time RT-PCR assays were used in this study: a modified version of the CDC N2 EUA assay [(Q)N2] and a second assay targeting the surface glycoprotein (GP). The (Q)N2 assay uses unmodified primers and probe (IDT); however, the assay itself was optimized for use with non-EUA reagents due to a world-wide reagent shortage. This assay was a 100% match to 99.62% of the SARS-CoV-2 sequences available (n = 3,423) in GenBank as of 22 May 2020. The GP assay, designed using the SARS-CoV-2 sequences publicly available as of 23 January 2020 (n = 7), targets a region of GP that differentiates SARS-CoV-2 from SARS-CoV. Subsequent analysis of all available SARS-CoV-2 genomes available at GenBank as of 22 May 2020 found an exact match to 99.56% of the available SARS-CoV-2 sequences (n = 3,644).

*Assay Optimization.* Primer and probe concentrations were optimized for use on the Roche LightCycler 480 (Roche Applied Science, Indianapolis, IN) using the SuperScript One-Step RT-PCR Kit (Thermo Fisher Scientific) and SARS-CoV-2 virus (UCC# R4714b, Unified Culture Collection, USAMRIID). Cycling conditions were: 50°C for 15 min; 95°C for 5 min; 45 cycles (95°C x 5 sec, 50°C x 30 sec, 60°C x 30 sec); and 40°C x 30 sec. Fluorescence readings were taken following each of the 45 cycles, and a sample was considered positive if the quantification cycle (Cq) was less than 40 cycles. Extracted nucleic acid was serially diluted in nuclease-free water and tested in triplicate with both assays to determine the preliminary limit of detection (LOD), the lowest concentration where all three replicates test positive. The preliminary LOD was then confirmed by testing 60 replicates at the preliminary LOD and increasing the concentration used until at least 58 of the 60 replicates test positive. Exclusivity testing was conducted using nucleic acid from 103 different organisms (46 DNA and 57 RNA viruses, bacteria, and parasites) including multiple respiratory pathogens and eight different coronaviruses.

*Assay performance.* Assay performance of the (Q)N2 assay in clinical matrix was determined using nonhuman primate nasopharyngeal (NP) swab matrix acquired from USAMRIID’s Veterinary Medicine Division using standard veterinary methods and viral transport media (VTM). Individual NHP NP samples were processed and tested for the presence of SARS-CoV-2 using the (Q)N2 real-time RT-PCR assay, and all negative NP samples were pooled. Live SARS-CoV-2 was serially diluted in buffer (PBS) and NP swab matrix. Total nucleic acid was extracted in triplicate [100 ml sample plus 300 ml TRIzol LS (ThermoFisher)] at each dilution using the Qiagen EZ1 Advanced XL Robot and the Virus RNA Mini Kit 2.0. Extracted nucleic acid was tested in duplicate using the (Q)N2 assay for a total of 6 replicates at each dilution. The preliminary LOD in NP matrix was confirmed by running 60 replicate extractions and testing each extraction in singlet.

*Screening pre- and post-challenge NHP NP swabs.* NP swabs were qualitatively analyzed for the presence of SARS-CoV2 genomic material using two different SARS-CoV-2 RT-PCR assays (see supporting methods). TRIzol LS –inactivated samples were extracted using QIAGEN QIAamp Viral RNA Mini Kit. Prior to extraction samples were spiked with High Concentration Internal Control (Qiagen), which was run in accordance with the manufacturer’s guidelines, to ensure the efficacy of the extraction procedure. Extracted samples were analyzed using the Applied Biosystems 7500 Fast Dx platform using primer/probe sets described above. Primers and probes used as described in with SuperScript II Platinum Taq. Each set of reactions was run with a SARS-CoV2 stock virus, extracted as a positive control, and a known negative NHP NP swab as a negative matrix control. A standard curve was generated and run alongside the N2 samples as an additional internal control for amplification. A standard curve was not available for the GP assay.

SARS-CoV-2 microneutralization assay

*Detection antibody screening/optimization:* Vero-E6 and Vero-76 cells, seeded in a 96-well plate, were inoculated with SARS-CoV-2/MT020880.1 at a MOI = 0.2 and incubated for 1 hr at 37C/5% CO2/80% humidity. Infection media was then removed and cells were washed once with 1X PBS, followed by addition of fresh cell culture media and incubation at 37C/5% CO2/80% humidity. Culture media was removed 24 hrs post-infection and cells were washed once with 1X PBS. PBS wash was removed and plates were submerged in formalin fixing solution for 24 hrs, then permeabilized with 0.2% Triton-X for 10 min at room temp and treated with blocking solution (3% BSA/PBS). Cells were incubated with primary antibodies (Sino Biological 40143-R0001; Invitrogen PA1-41098; Novus Biologicals NB100-56048 and NB100-56578), diluted 1:500 in blocking solution, for 2 hrs at room temperature (RT). Cells were washed prior to addition of Alexa Fluor 488 conjugated secondary antibody diluted 1:2000 in blocking solution for 1 hr at RT, then washed and counterstained with DAPI.

*Multiplicity of infection optimization:* Vero-E6 cells, seeded in a 96-well plate, were inoculated with serially diluted SARS-CoV-2/MT020880.1 at MOIs between 0.6 and 0.007 and incubated for 1 hr at 37C/5% CO2/80% humidity. Infection media was then removed and cells were washed once with 1X PBS, followed by addition of fresh cell culture media and incubation at 37C/5% CO2/80% humidity. Culture media was removed 24 or 48 hrs post-infection and cells were washed once with 1X PBS. PBS wash was removed and plates were submerged in formalin fixing solution for 24 hrs, then permeabilized with 0.2% Triton-X for 10 min at room temp and treated with blocking solution (3% BSA/PBS). Infected cells were detected using a primary detection antibody recognizing the SARS-CoV-2 nucleocapsid protein (Sino Biological, 40143-R001) and Alexa Fluor 488 conjugated secondary antibody (goat α rabbit), then counterstained with DAPI, as described above. Infected cells were enumerated using Operetta high content imaging instrument and data analysis was performed using the Harmony software (Perkin Elmer).

*Microneutralization Assay.* A pre-titrated amount of authentic SARS-CoV-2/MT020880.1 virus at final multiplicity of infection of 0.2 was incubated with serial dilutions of heat inactivated serum for 1 hr at room temperature. The antibody-virus mixture was applied to monolayers of Vero-E6 cells in a 96-well plate and incubated for 1 hr at 37°C in a humidified incubator. Infection media was then removed and cells were washed once with 1X PBS, followed by addition of fresh cell culture media. Culture media was removed 24 hrs post-infection and cells were washed once with 1X PBS. PBS wash was removed and plates were submerged in formalin fixing solution for 24 hrs, then permeabilized with 0.2% Triton-X for 10 minutes at RT and treated with blocking solution (3% BSA/PBS). Infected cells were detected using a primary detection antibody recognizing the SARS-CoV-2 nucleocapsid protein (Sino Biological, 40143-R001) and Alexa Fluor 488 conjugated secondary antibody (goat α rabbit), then counterstained with DAPI, as described above. Infected cells were enumerated using Operetta high content imaging instrument and data analysis was performed using the Harmony software (Perkin Elmer).

SARS-CoV-2 PRNT

An equal volume of complete media (EMEM+10% heat inactivated FBS+1% Pen/Strep, 0.1% gentamycin, 0.2% fungizone) containing virus (SARS-CoV-2 Washington isolate) is combined with 2-fold serial dilutions of complete media containing heat inactivated serum and incubated at 37°C in a 5% CO_2_ incubator for 1 hr (total volume 222 microliters). 180 ul per well of the combined virus/antibody mixture is then added to VERO76 ATCC cell monolayers in 6-well plates and allowed to adsorb for 1 hour at 37°C. 3 mL per well of agarose overlay (0.6% SeaKem ME agarose, EBME with HEPES, 10% heat inactivated FBS, 100X nonessential amino acids, and antibiotics) is then added and allowed to solidify at room temperature. The plates are placed in a 37°C incubator for 2 days and then 2 mL per well of agarose overlay containing 5% neutral red and 5% heat inactivated FBS is added and the plates are returned to 37°C. After 1 day the plaques are counted on a light box. The PRNT50 and PRNT80 titers are the reciprocal of the highest dilution that results in a 50% or 80% reduction in the number of plaques relative to virus in the presence of complete media (no antibody).

Euroimmun SARS-CoV-2 S1 ELISA

Serum samples were screened with the Euroimmun SARS-CoV-2 S1 ELISA (Euroimmun, EI 2606-9601 G) as per kit instructions. Briefly, the kit materials were brought to room temperature for 30 minutes. Serum samples were diluted 1:101 using the supplied sample buffer. 100µL of the diluted samples, supplied controls, and supplied calibrator were added to the pre-coated wells and incubated at 37°C for 1 hour. After 1 hour, the plate was washed 3 times with 300uL of supplied wash buffer using a microplate washer (Biotek 405TS). 100µL of enzyme conjugate was added to the wells and incubated at 37°C for 30 minutes. The plate was washed 3 times as outlined above prior to adding 100µL of substrate for 30 minutes at RT. Finally, 100µL of stop solution was added prior to reading absorbance at 450nm, with a reference wavelength at 635nm (Tecan M200). Data was processed according to kit instructions to determine negative, positive, or borderline results.

SARS-CoV-2 MAGPIX Multiplex Immunoassay

*Magnetic microsphere production:* Recombinant SARS-CoV-2 full trimeric spike (gift from Dr. Jason McLellan’s group; UT-Austin^1^), S1 (Sino Biological, 40591-V08H), RBD (Sino Biological, 40592-V08H), NP (Native Antigen Company, REC31812-100), and SARS-CoV-1 full spike (Protein Sciences, no catalogue number) proteins were conjugated to magnetic microspheres using the Luminex xMAP® antibody coupling kit (Luminex Inc., Austin, TX, USA) according to the manufacturer’s instructions. Briefly, 500 µL of Magplex microspheres (12.5 × 106 microspheres/mL) were washed three times using a magnetic microcentrifuge tube holder and resuspended with 480 µL of activation buffer. Then, 10 µL of both sulfo-NHS and EDC solutions were added to the resuspended microspheres. The tube was covered with aluminum foil and placed on a benchtop rotating mixer for 20 min. After surface activation with EDC, the microspheres were washed three times with activation buffer prior to adding the recombinant protein antigen at a final concentration of 4 µg antigen/1 × 10^6^ microspheres. This concentration of recombinant protein coupled to the surface of microspheres has shown to be optimal for IgG and IgM detection^2^. The tube was again covered with aluminum foil and placed on a benchtop rotating mixer for 2 hr. After this coupling step, the microspheres were washed three times with wash buffer and resuspended in 500µL of wash buffer for further use. CoV2 full spike, S1, RBD, NP, and CoV1 full spike were coupled to Magplex microsphere regions #45, #55, #65, #25, and #77 (Luminex Inc., Austin, TX, USA), respectively, in order to facilitate multiplexing experiments. Beads were stored at 4°C until further use.

*Screening pre- and post-challenge NHP serum.* Serum samples were diluted at 1:100 phosphate buffer saline (PBS) with 0.02% Tween-20 (PBST) with 5% skim milk (PBST-SK). Serial dilutions were made a 1/3 dilutions starting at 1:900. Each individual antigen-coupled bead were mixed at a 1:1 ratio prior to diluting in PBST to 5 × 10^4^ microspheres/mL and added to the wells of a Costar polystyrene 96-well plate at 50 µL per well (2500 microspheres of each antigen bead set/well). The plate was placed on a magnetic plate separator (Luminex Inc.) covered with foil, and microspheres were allowed to collect for 60 sec. While still attached to the magnet, the buffer was removed from the plate by inverting and disposing into the sink (for pre-bleed serum samples in BSL2) or into a kill plan with 5% Microchem (for post-challenge samples in BSL3) . Then, 50 µL of diluted serum samples were added to appropriate wells. The plate was covered with a black, vinyl plate cover and incubated with shaking for 1 hr at RT. The plate was washed three times with 100 µL of PBST for each wash, using the plate magnet to retain the Magplex microspheres in the wells. Liquid was discarded as above for BSL2 or BSL3 sample processing. 50 µL of a 1:100 dilution of mouse anti-human IgM phycoerythrin conjugate (Invitrogen, MA1-10381) or goat anti-human IgG phycoerythrin conjugate (Sigma, P9170) in PBST-SK was added to the wells. The plate was covered again with a black, vinyl plate sealer and incubated with shaking for 1 hr at RT. After incubation, the plate was washed three times as detailed above and the Magplex microspheres were resuspended in 100 µL of PBST for analysis on the Magpix instrument. Raw data was reported as median fluorescence intensity for each bead set in the multiplex.

Genomic Analysis

*Library preparation – Stock Characterization by ARTIC primer amplification.* cDNA synthesis was performed with the Superscript IV first-strand synthesis system (Life Technologies/Invitrogen, Carlsbad, CA). Multiplex PCR was performed with the ARTIC primer set, which was designed to amplify overlapping regions of the Sars-CoV-2 reference genome (MN908947.3). Primer information and genomic alignment position is available here: https://github.com/artic-network/artic-ncov2019/tree/master/primer_schemes/nCoV-2019/V1. PCR products were purified with the MinElute PCR purification kit (QIAgen, Valencia, CA). Libraries were prepared with the SMARTer PrepX DNA Library Kit (Takara Bio, Mountain View, CA), using the Apollo library prep system (Takara Bio, Mountain View, CA). The libraries were evaluated for quality using the Agilent 2200 TapeStation (Agilent, Santa Clara, CA). After quantification by real-time PCR with the KAPA SYBR FAST qPCR Kit (Roche, Pleasanton, CA), libraries were diluted to 10 nM.

*Sequencing.* The chosen platform for re-sequencing for this study was the Illumina MiSeq. For more information about the Illumina sequencer, follow the link to the Illumina website (http://www.illumina.com/publications/overview.ilmn).

*Sequence analysis*. Raw sequences from the Illumina MiSeq were cleaned and aligned using an in-house perl wrapper, VSALIGN, developed by CGS. VSALIGN performs read cleaning and quality control using PRINSEQ-lite and in-house scripts. This cleaning and pre-processing step included adapter removal, duplicate/chimeric read removal, and quality filtering, keeping reads with an average quality score > 25 (phred). Pre-processed and filtered reads were aligned to the strain MT020880.1, which the USAMRIID stock is derived from, using DNAStar Lasergene nGen. Changes relative to the MT020880.1 sequence required greater than 50% of the reads supporting the change, with at least 20x coverage.

*Phylogenetics.* SARS-CoV-2 sequences were aligned with MAFFT v7.220 using default settings followed by manual curation using Geneious version R9. The statistical selection of the best-fit nucleotide substitution model was performed with jModelTest2. We reconstructed a Maximum-likelihood tree based on the GTR+Γ4 model using the software FastTree v2.1 (http://www.microbesonline.org/fasttree/ ) under an exhaustive search, and computing 5000 local support values (Shimodaira-Hasegawa test).

USAMRIID ABSL PPE Stance and Discovery of NP Swab Positive NHP

Comparative Medicine Division implemented a respiratory protection equipment requirement of N-95 or above for ABSL2 rooms, in early February 2020 as an added protective measure against latent TB activation in the colony. USAMRIID screens all personnel that could interact with NHPs for TB biannually, as it is known that NHPs are susceptible to TB transmission from humans. However, the institute decided to take additional protective measures to further control the integrity of the colony. On February 26th, The Comparative Medicine Division switched to an ABSL3 PPE posture (PAPR, Tyvek, double gloves with bleach decon) due to a lack of N-95s. Due to COVID19 concerns, on 13 March, several changes were made to operations. Personnel were instructed to put on PPE prior to going in to the NHP corridors (rather than in anterooms). All items going in to the animal corridors were deconned prior to movement in. All NHP cages were cleaned in place to minimize movement of animal caging between areas. All personnel with offices in NHP housing areas were relocated and social distancing was implemented shortly thereafter. On April 6th, a series of nonhuman primate nasopharyngeal swabs were acquired from USAMRIID nonhuman primates for use for baselining in a SARS-CoV-2 real-time PCR assay qualification study by the Diagnostic Systems Division. Samples were taken from 12 different animals (4 African Greens, 4 Rhesus macaques, 4 Cynomolgus macaques). All 12 animals tested were subsequently moved to ABSL3 for study use on April 13th. Upon discovering the positive swab in one of the African Greens, the room from which this animal was removed was placed on quarantine the morning of the April 16th. There were no animals reported for in appetence or respiratory signs.

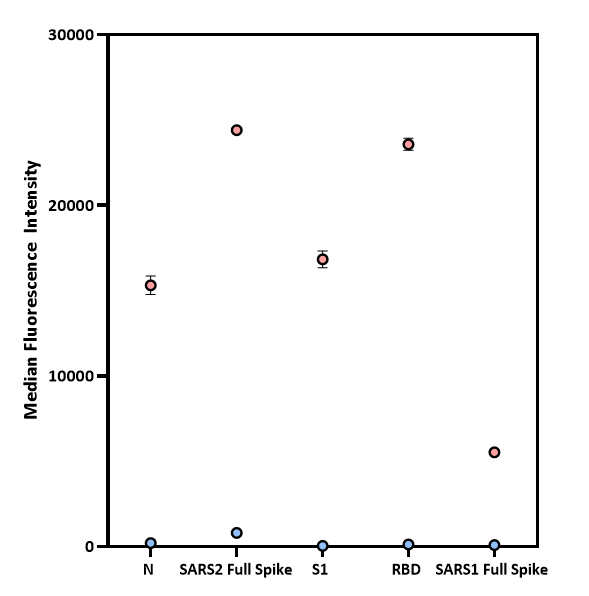

**Fig S1.** Verification of coupling efficacy of Magplex microspheres using a SARS-CoV-2 positive and negative control. All samples were run in duplicate.

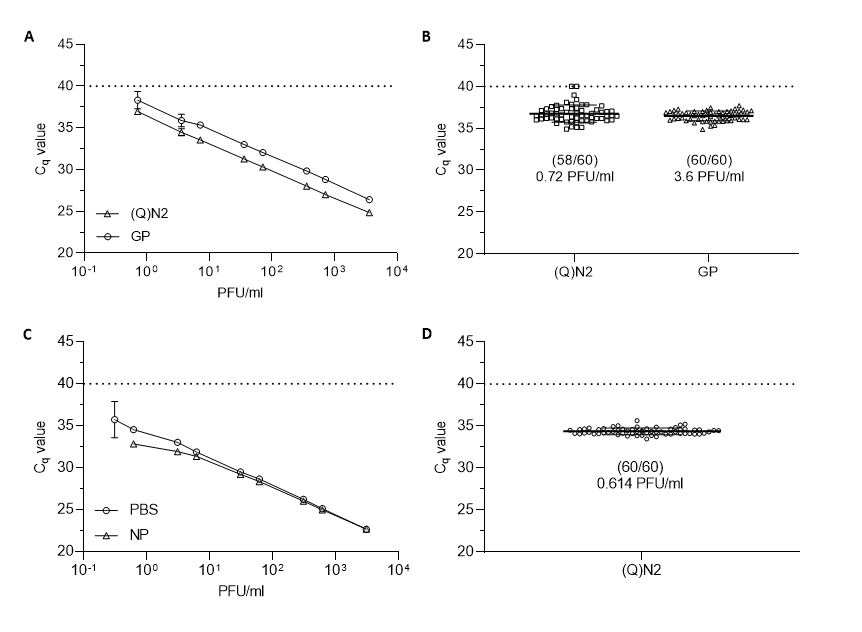

**Fig S2 . Characterization of assay performance.** (A) Extracted SARS-CoV-2 nucleic acid was run in triplicate with the (Q)N2 and GP assays with the preliminary LOD being 0.72 PFU/ml. (B) Confirmation of LOD testing resulted in 58 of 60 replicates testing positive with the (Q)N2 assay at the preliminary LOD while the GP assay required increasing the nucleic acid concentration to 3.6 PFU/ml. Live virus was serially diluted in NP swab matrix and PBS (C)and tested with the (Q)N2 assay in duplicate=. Sixty replicates of live virus diluted in NP swab (D) at the preliminary LOD were extracted and tested with the (Q)N2 assay, increasing the amount of virus used until 58 of the 60 replicates tested positive. All error bars are the standard deviation.

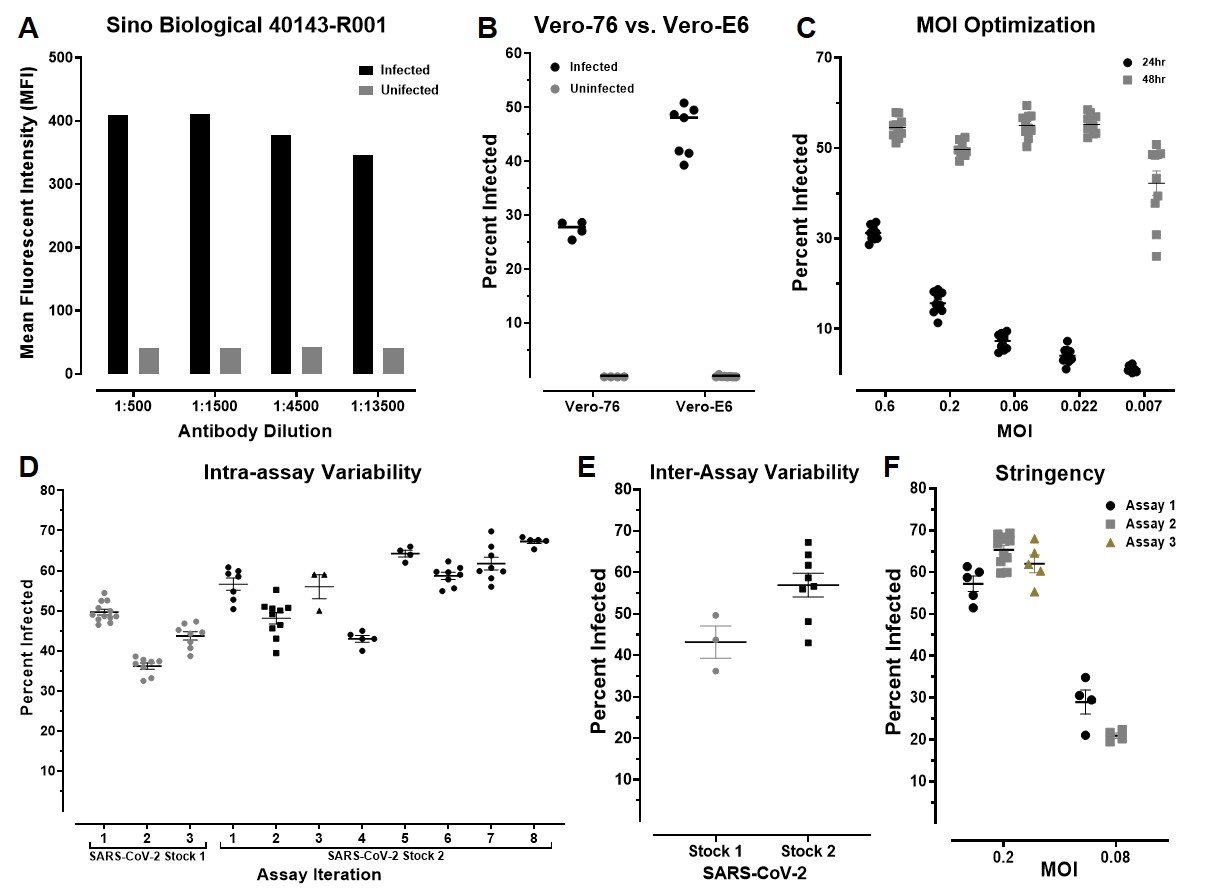

**Fig S3. SARS-CoV-2 microneutralization assay development.** (A) SARV-CoV-2 infected Vero-76 cells were immuno-stained with indicated concentration of primary antibody 40143-R001 and Alexa Fluor 488 conjugated secondary antibody. Mean fluorescent intensity of infected and uninfected cells was determined to optimize signal-to-noise ratio. (B) Vero-76 and Vero-E6 cells were infected with SARS-CoV-2 and an MOI of 0.4 for 24 hours. Fixed cells were immuno-stained and the percent of infected cells was determined. (C) Vero-E6 cells were infected with SARS-CoV-2 at indicated MOI for 24 or 48 hours. Fixed cells were immuno-stained to determine percent of infected cells. (D) Vero-E6 cells were infected with two stocks of SARS-CoV-2 at MOI 0.4 for 24 hours and immuno-stained to determine percent of infected cells across individual assay iterations. (E) Inter-assay variability of experiments described in D was determined for each stock. Individual data points represent the average percent infection from each plate within an iteration. (F) Vero-E6 cells were infected with SARS-CoV-2 an indicated MOI for 24 hours and immuno-stained to determine percent of infected cells across individual assays. (B & C) Individual data points represent the percent of infected cells from replicate virus only treated wells. (D & F) Individual data points represent the average percent infection of at least 4 replicate virus only treated wells from the same assay plate.

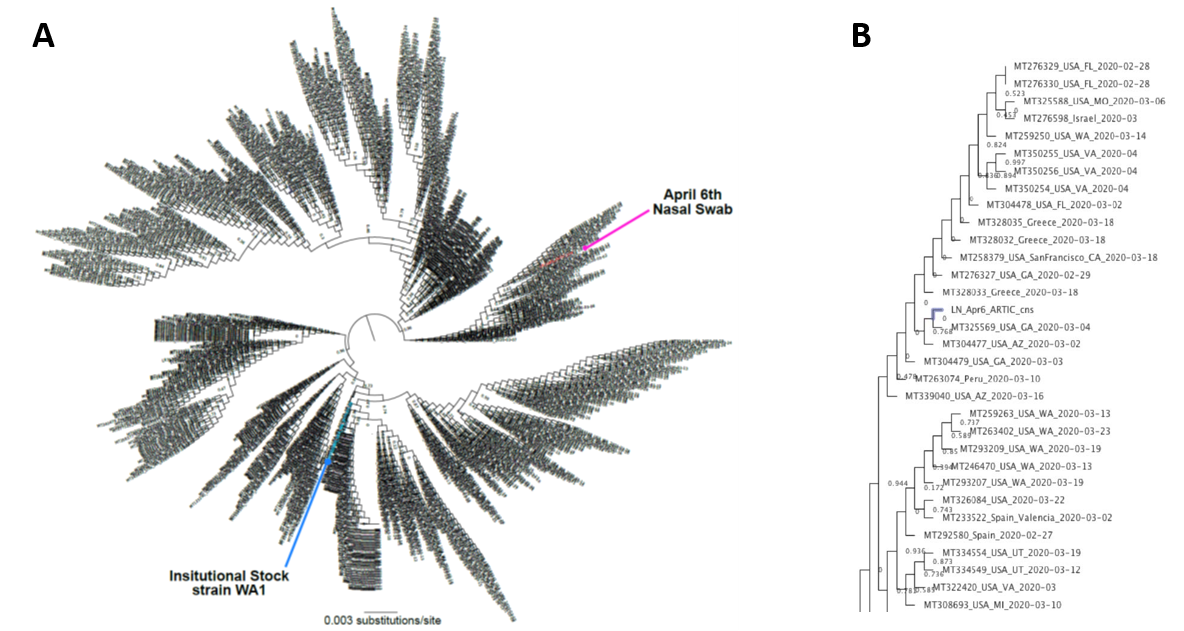

**Fig S4. Phylogenetic Summary of AGM1 Pre-Screen NP swab**. A) Radial phylogram. B) Expanded APR6 Tree. A radial Maximum-likelihood tree estimated using 1,003 SARS-CoV-2 coding-complete genomes. The WA1 strain (Genbank accession number: MT020880) and consensus genome sequence from an infected non-human primate nasal swab collected on April 6th are highlighted using colored circles. Tree branches are scaled by substitutions per site. Tree node support values were generated using 5000 Shimodaira-Hasegawa tests and are shown in decimal form.

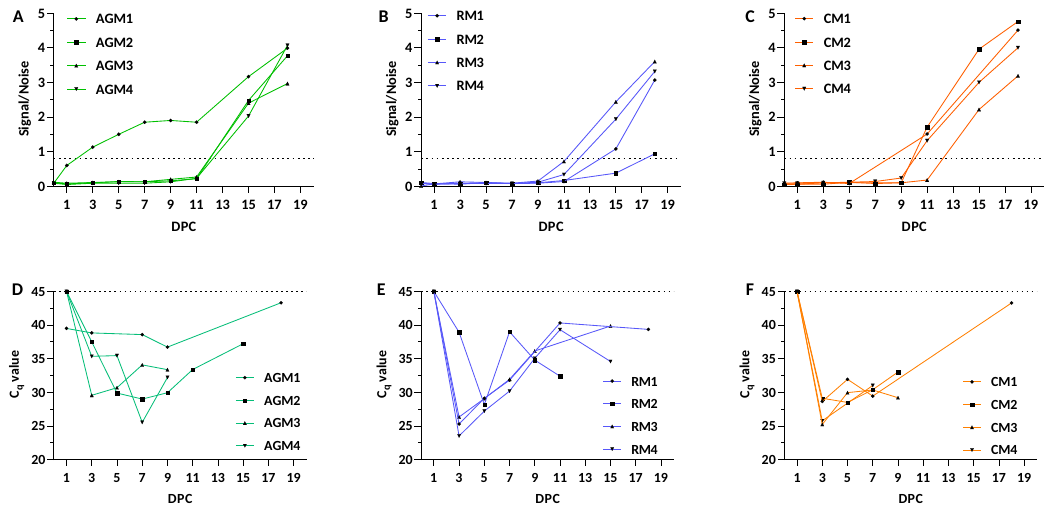

**Fig S5. Longitudinal Immune and Viral Response.** Longitudinal response for AGM (A and D), RM (B and E), and CM (C and F) cohorts for IgG in serum samples and viral load in NP swabs. Longitudinal samples were collected on day 1 and every two days thereafter. Subsequent sampling was done on days 15 and 18. Only the PCR positive samples are shown in graphs D-F. Serum samples were screened by commercial Euroimmun SARS-CoV-2 S1 IgG ELISA kit. This direct ELISA kit measures IgG response against the S1 subunit of the spike glycoprotein.

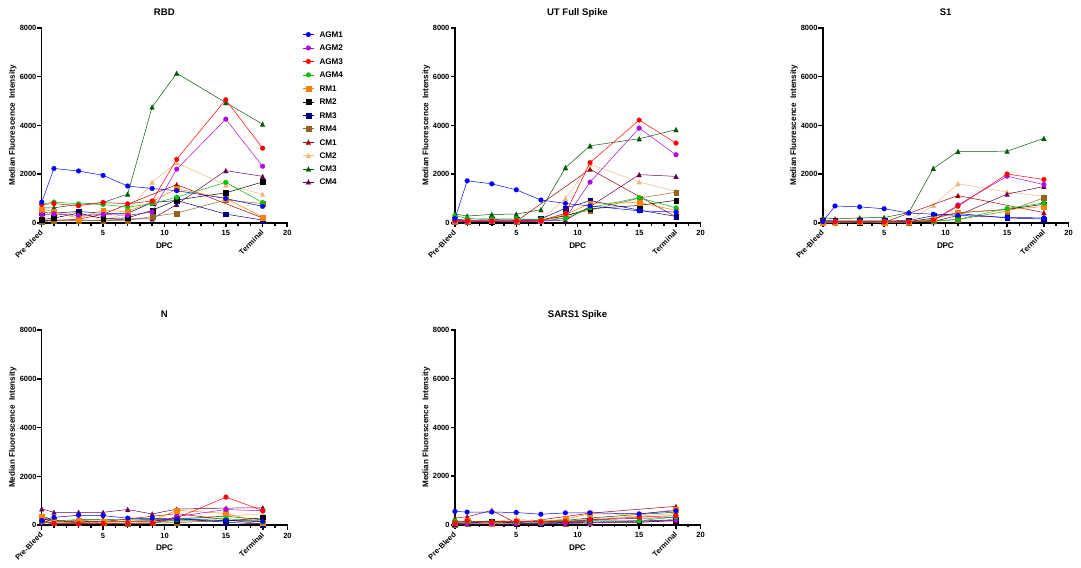

**Fig S6.** Longitudinal IgM response for Magpix multiplex. All samples were run in duplicate at each time point.

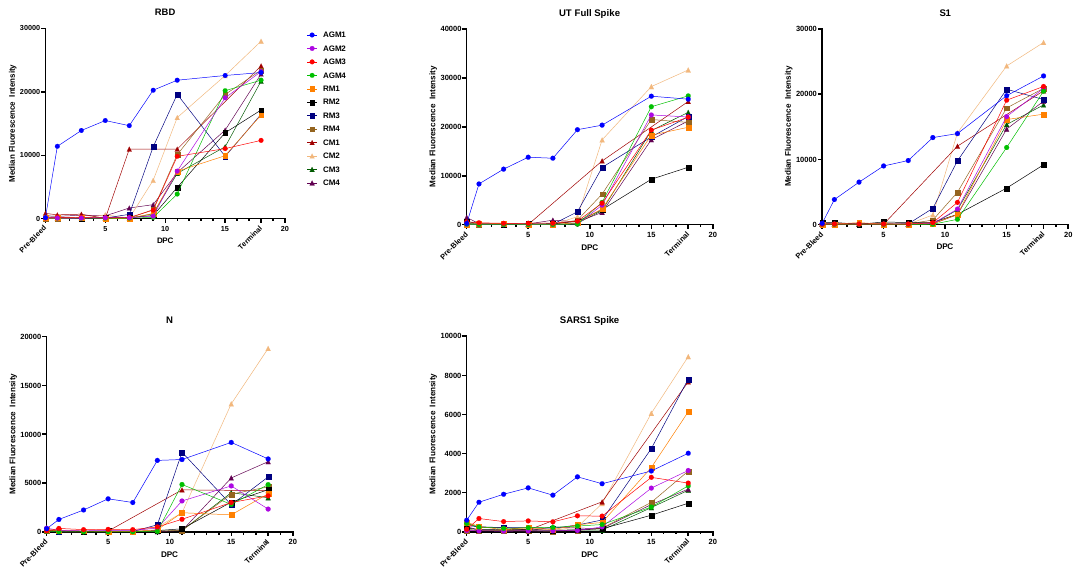

**Fig S7.** Longitudinal IgG response for Magpix multiplex. All samples were run in duplicate at each time point.

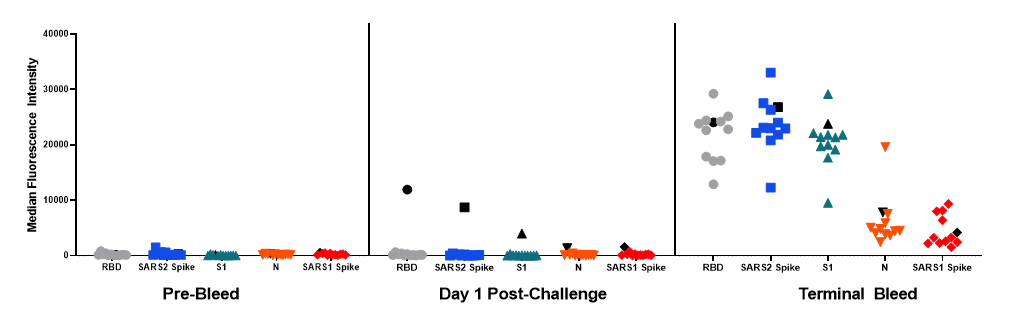

**Fig S8. Comparison of IgG response.** Pre-bleed, in-life, and terminal bleed NHP serum were screened by the COVID Magpix multiplex immunoassay for IgG response to five different targets: Full trimeric SARS-CoV-2 spike, receptor binding domain (RBD), S1 subunit, nucleoprotein (N), and full SARS-CoV-1 spike. The pre-exposed NHP is represented by black data points in each graph. All NHPs were IgG negative on the date of the pre-bleed. On day 1 post-challenge, all NHPs were negative with the exception of the pre-exposed NHP. By the terminal bleed, all NHPs had measurable IgG response to the multiplex panel. The pre-exposed NHP was not statistically different from the average response of all the NHPs for the pre and terminal bleeds; however, all five assays were statistically significant for the pre-exposed NHP at day 1 post-challenge. Statistical significance determined using the Holm-Sidak method, with alpha = 0.05.

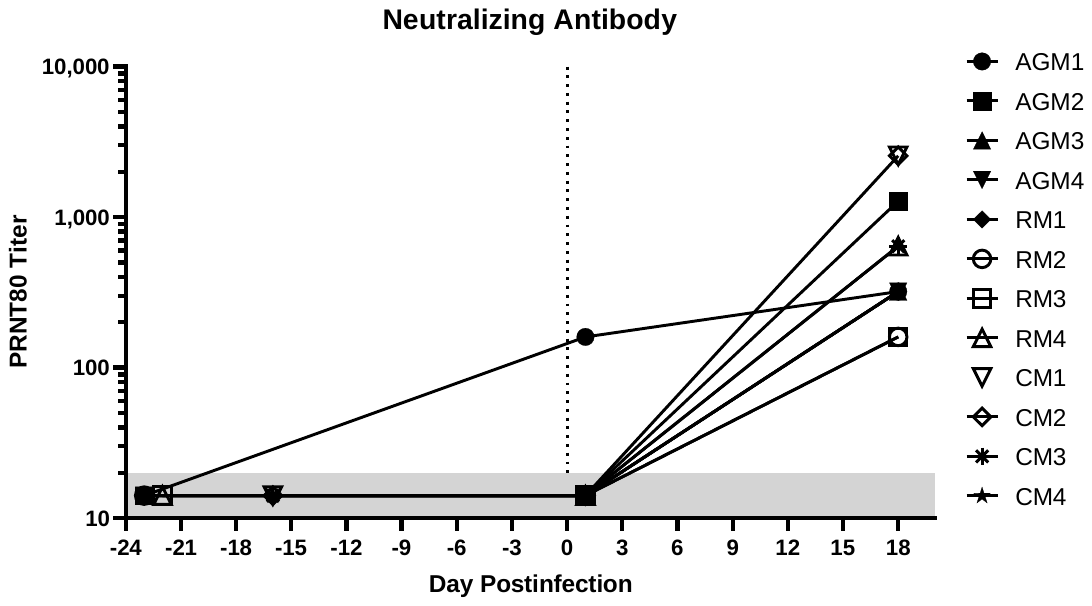

**Fig S9.** PRNT80 Titer at pre-bleed, D1, and terminal time points.

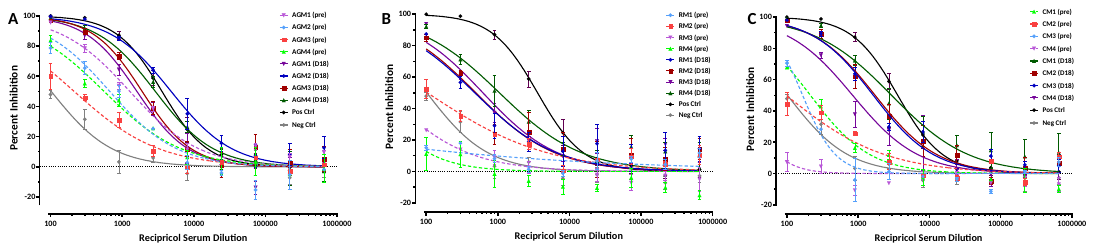
 **Fig S10.** Serially diluted serum for A) AGM, B) RM, and C) CM (pre-challenge and day 18 post-challenge), mixed with pre-tittered SARS-CoV-2, was added to Vero-E6 cells at an MOI of 0.2. Cells were fixed 24 hours post-inoculation and immuno-stained with virus specific antibody to enumerate the number of infected cells by fluorescent microscopy (Operetta high-content imaging). Percent of infected cells was determined using Harmony software and percent inhibition calculated relative to untreated cells. Data is derived from a minimum of 2 technical replicates and from at least 2 replicate assays.

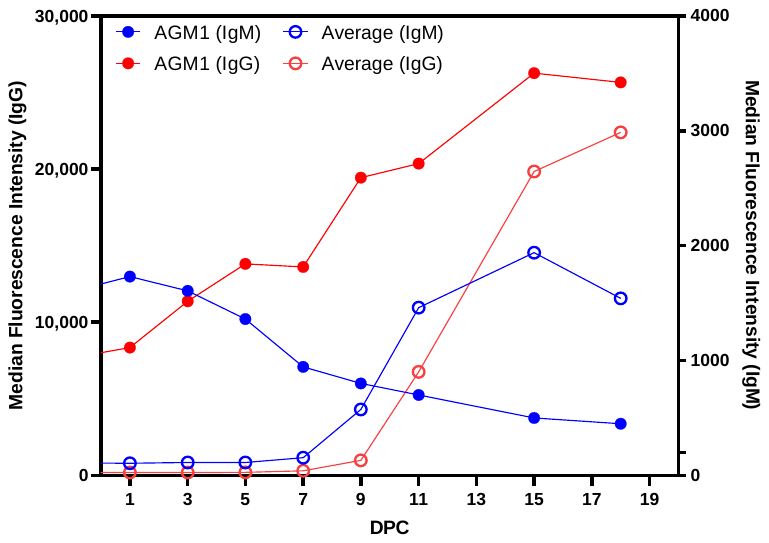

**Fig S11.** Comparison of longitudinal IgM and IgG response to SARS-CoV-2 full spike protein of the average NHP cohort versus the pre-exposed AGM1.

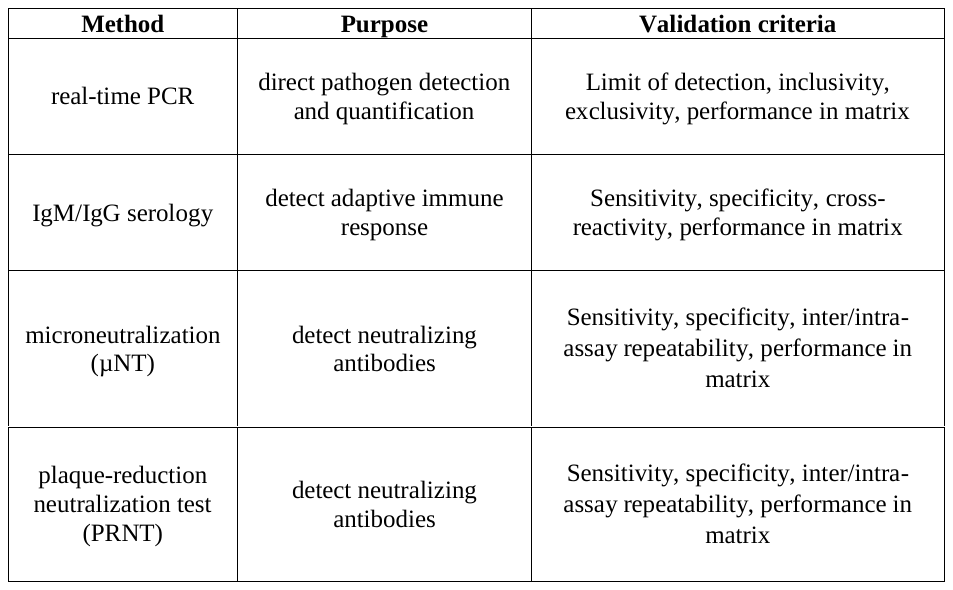

**Table S1.** Assays for SARS-CoV-2 animal model and countermeasure development

| **General Identifiers** | | **Immunoassay Data** | | | **Neutralization Data** | | **PCR Data** | | |
| --- | --- | --- | --- | --- | --- | --- | --- | --- | --- |
| **Animal ID** | **Species** | **EuroImmun ELISA** | **Magpix S1 Assay** | **Result** | **Heat Inactivated PRNT**_50_ | **Result** | **(Q)N2** | **GP** | **Result** |
| **AGM1** | African Green | 0.02 | 161 | **NEG** | <20 | **NEG** | 28.99 | 31.2 | **POS** |
| **AGM2** | African Green | 0.02 | 60 | **NEG** | <20 | **NEG** | ND | ND | **NEG** |
| **AGM3** | African Green | 0.03 | 46 | **NEG** | <20 | **NEG** | ND | ND | **NEG** |
| **AGM4** | African Green | 0.02 | 42 | **NEG** | <20 | **NEG** | ND | ND | **NEG** |
| **CM1** | Cyno | 0.03 | 83 | **NEG** | <20 | **NEG** | ND | ND | **NEG** |
| **CM2** | Cyno | 0.02 | 36 | **NEG** | <20 | **NEG** | ND | ND | **NEG** |
| **CM3** | Cyno | 0.02 | 45 | **NEG** | <20 | **NEG** | ND | ND | **NEG** |
| **CM4** | Cyno | 0.03 | 50 | **NEG** | <20 | **NEG** | ND | ND | NEG |
| **RM1** | Rhesus | 0.02 | 41 | **NEG** | <20 | **NEG** | ND | ND | **NEG** |
| **RM2** | Rhesus | 0.02 | 254 | **NEG** | <20 | **NEG** | ND | ND | **NEG** |
| **RM3** | Rhesus | 0.02 | 51 | **NEG** | <20 | **NEG** | ND | ND | **NEG** |
| **RM4** | Rhesus | 0.02 | 113 | **NEG** | <20 | **NEG** | ND | ND | **NEG** |

**Table S2.** Summary of pre-screening data.

| **Sample** | **Test** | **(Q)N2** | | **GP** | |
| --- | --- | --- | --- | --- | --- |
|  |  | **positive (total)** | **AVE Cq** | **positive (total)** | **AVE Cq** |
| NP swab (left) | initial | 3 (3) | 28.99 | 3 (3) | 31.2 |
| NP swab (right) | initial | 0 (3) | - | - | - |
| NP swab (left) | repeat | 9 (9) | 29.2 | 9 (9) | 31.3 |
| NP swab (right) | repeat | 0 (9) | - | 4 (9) | 35.3 |

**Table S3.** NHP NP swab PCR sample testing results from the initial pre-screen swab and the D1 post-challenge swab for AGM1.

| **NHP ID** | **IC50 (Recipricol Dilution)** | | **Fold Change** |
| --- | --- | --- | --- |
|  | **D 1** | **D 18** |  |
| **AGM1** | 1274 | 1603 | 1.26 |
| **AGM3** | 212.8 | 1894 | 8.90 |
| **AGM4** | 557.6 | 2895 | 5.19 |
| **AGM2** | 668.4 | 4812 | 7.20 |
| **RM1** | 5 | 476.9 | 95.38 |
| **RM2** | 103.7 | 493.3 | 4.76 |
| **RM3** | 25.8 | 619.1 | 24.00 |
| **RM4** | 15.65 | 1053 | 67.28 |
| **CM1** | 213.6 | 2939 | 13.76 |
| **CM2** | 91.32 | 1730 | 18.94 |
| **CM3** | 166.2 | 1631 | 9.81 |
| **CM4** | 24.96 | 853.7 | 34.20 |
| **Neg Ctrl** | 101.8 | | n/a |
| **Pos Ctrl** | 3589 | | n/a |

**Table S4.** Microneutralization Calculated IC50 for D1 and D18 post-challenge.
